# The proton-activated receptor TDAG8 is upregulated in oligodendrocytes during maturation and under acidic conditions

**DOI:** 10.1101/2023.03.02.530787

**Authors:** Mikołaj Opiełka, Fionä Caratis, Martin Hausmann, Maria Velasco-Estevez, Cheryl de Vallière, Klaus Seuwen, Gerhard Rogler, Bartosz Karaszewski, Aleksandra Rutkowska

## Abstract

Acidosis is one of the hallmarks of demyelinating central nervous system (CNS) lesions in multiple sclerosis (MS). Response to acidic pH is primarily mediated by a family of G protein-coupled proton-sensing receptors: OGR1, GPR4, and TDAG8. These receptors are inactive at alkaline pH, while at acidic pH they are maximally activated. Genome-wide association studies identified a locus within the TDAG8 gene to be associated with several autoimmune diseases including MS. Notably, we here found that TDAG8 expression is upregulated in MS plaques which prompted us to explore the expression and function of TDAG8 in the CNS in human MO3.13 oligodendrocytes *in vitro* and *in vivo* in the lipopolysaccharide-induced neuroinflammation model. We found that TDAG8 is upregulated in maturing oligodendrocytes and temporarily under acidic conditions. Acidic pH also induces oligodendrocyte branching, inhibits chemotaxis and affects the expression of oligodendrocyte maturation markers, PDGFRα and MBP *in vitro*. Even though myelination was not affected in the adult TDAG8-deficient mice, the expression of human and murine TDAG8 was strongly regulated upon inflammation *in vivo* in the brain and *in vitro* in lipopolysaccharide and pro-inflammatory cytokine-treated oligodendrocytes. Together these findings point toward a potential role of TDAG8 in oligodendrocyte biology, neuroinflammation and pathophysiology of MS and provide new directions for further scientific enquiry.

## INTRODUCTION

To maintain pH homeostasis, cells are required to sense acidic changes in their environment and respond accordingly. Three G protein-coupled receptors (GPCRs), the ovarian cancer GPCR 1 (OGR1, GPR68), GPCR 4 (GPR4) and T cell death-associated gene 8 (TDAG8, GPR65) have been shown to sense extracellular protons and stimulate differing signalling pathways [1,2]. The receptors are activated by acidic extracellular pH, through the protonation of several histidine residues located on the extracellular surface of the receptor [2]. At pH 7.6-7.8, the receptors are almost silent, and at pH 6.8-6.6, they are maximally activated [2].

Upon extracellular acidification, which is frequently observed in inflammation, proton activation of TDAG8 has been linked with anti-inflammatory roles in various disease models and in cells including inhibition of granulocyte, macrophage and monocyte inflammatory processes [3,4]. In an earlier study, we examined the role of TDAG8 in inflammation in the experimental colitis, dextran sodium sulfate (DSS), model. We reported that TDAG8-deficiency leads to increased macrophage and neutrophil infiltration, and increased expression of pro-inflammatory mediators in both, the acute and chronic DSS models of colitis. Further, we examined acidosis-activated and TDAG8-mediated pathways in peritoneal macrophages by RNA sequencing. We found that activation of TDAG8 by low pH in peritoneal macrophages modulated the expression of genes involved in the immune response [5]. Additionally, as was demonstrated by us and others, TDAG8 can promote changes in cell adhesion and contractility, gene expression, cell division and proliferation [6].

Neuroinflammation, neurodegeneration, ischemic stroke and ageing are associated with acidosis in the central nervous system (CNS) [7–9]. In mice subjected to the middle cerebral artery occlusion (MCAO), the pH measured in the infarct region decreases to 6.5 compared to the ipsilateral hemisphere [10]. In Alzheimer’s disease, a neurodegenerative disease of the CNS, lower pH was found in the cerebrospinal fluid (CSF) and post-mortem brains [7]. Similarly, in multiple sclerosis (MS), a chronic autoinflammatory disease of the CNS, increased lactate levels were reported in the affected brain tissue [8]. Also in animals, a decreased pH was demonstrated in the spinal cords of mice in the experimental autoimmune encephalomyelitis (EAE) model of MS [11]. In the same mouse model, but in a different study, CNS acidosis was shown to be associated with demyelination and oligodendrocyte injury [12].

Despite the identification of TDAG8 as a risk gene for MS in high-powered genome-wide association studies (GWAS) [13,14], limited research has been reported in this area. Only a couple of studies investigated the expression and signalling of TDAG8 in the CNS to date. For instance, expression of TDAG8 was reported in circumventricular organs in mouse microglia [15]. In this study, TDAG8-mediated signalling was involved in CO_2_-evoked freezing and sympathetic reactions in a mouse model of panic disorder. The TDAG8-deficient mice exhibited reduced activation of microglial cells and IL1β release in the subfornical area [15]. Additionally, it was demonstrated that the IL1β release is severely attenuated in LPS-activated microglia in an acidic environment through a TDAG8-dependent mechanism [16]. The expression of TDAG8 in another CNS-specific cell type was investigated by Bortell and colleagues (2017). They observed that TDAG8 is one of the three strongest expressed genes in astrocytes after *in vitro* treatment with methamphetamine. Interestingly, the other two upregulated genes, MAP2K5 and CXCL5, similar to TDAG8, are also inflammation and immune regulators. In the CNS, TDAG8 was also found to be expressed in neurons and to be negatively involved in chronic inflammatory pain [18,19].

As for the role of TDAG8 in the EAE model, a single-cell RNA-sequencing (scRNA-seq) study and computational analysis used to determine different cellular states of Th17 cells revealed that TDAG8 promotes Th17 cell pathogenicity in the EAE model [20] via the Th17 cell master transcription factor, Rorγt, which binds the promoter region of TDAG8 [21]. However, another study where the effects of TDAG8 deficiency in the EAE model [22] were studied, found that TDAG8 deficient mice exhibit an exacerbated course of EAE and that TDAG8 signalling in invariant NKT cells, but not in CD4^+^ T cells, was responsible for attenuating the autoimmune responses [22]. Strikingly, the immunoregulatory effects of TDGA8 depended mostly on signalling in invariant natural killer (NK) cells, but not CD4+ T cells [22]. Yet another study reported protection from the EAE in mice with TDAG8-deficient T cells [20]. The involvement of TDAG8 signalling in psychosine-mediated oligodendrocyte death in Krabbe disease (globoid cell leukodystrophy), a genetic demyelinating disease caused by the accumulation of psychosine, was investigated by Giri et al. (2006). In this study, using the MO3.13 human oligodendrocytes, the authors showed that the toxic effects of psychosine (galactosylsphingosine) on oligodendrocytes are TDAG8-independent, also demonstrating that psychosine is not a TDAG8 ligand.

The effects of extracellular pH on oligodendrocyte function in a receptor-independent setting were investigated in the context of remyelination and MS [24]. Primary rat oligodendrocyte progenitor cells (OPCs) were cultured in media with pH ranging from 6.0 to 8.0 and cell adhesion, survival, migration, proliferation and differentiation were examined *in vitro*. The study reported reduced chemotaxis and velocity of rat OPCs in acidic pH and a tendency to migrate towards the low pH in a Zigmond chamber where a pH gradient from 6.0 to 7.0 was formed. The adhesion and length of OPCs increased in acidic pH, in line with the decreased migration reported in the low pH experiments. Moreover, the rat OPC viability, proliferation and differentiation were significantly decreased in the low pH of 6.0-6.5 compared to the neutral pH of 7.0-7.5.

Here, we broadly explored the expression and function of the proton-activated receptor TDAG8 in the mouse and human healthy and inflamed CNS to provide direction for future, more directed studies. The changes in TDAG8 expression in human MO3.13 oligodendrocytes, as well as the pH effects on cell migration, morphology and differentiation are examined.

## METHODS

### Human MS and healthy brains

Post-mortem frozen human brain sections from MS and control donors were obtained from the Rocky Mountain MS Center Tissue Bank (Englewood, CO, USA) after approval by the Medical University of Gdansk (Poland) bioethics committee (NKBBN/253/2018). Lightly frozen 1 cm by 1 cm fragments were dissected from the white matter (WM) and periventricular WM plaques. Corresponding areas of WM were dissected from control brains. The fragments were deep-frozen in liquid nitrogen, ground using mortar and pestle, and homogenized in Fenozol reagent (A&A Biotechnology, 203-50). Total RNA was isolated from approximately 200 mg of homogenized tissue using the Total RNA Mini kit (A&A Biotechnology, 031-100) in an RNAase-free environment according to the manufacturer’s instructions. RNA quantification and quality control were performed spectrophotometrically at 260 nm and 280 nm using a PerkinElmer VICTOR Nivo plate reader (Perkin Elmer, USA) equipped with a μDrop Plate (Thermo Scientific, N12391). Isolated total RNA was stored at -20°C until used.

### Animals

The lipopolysaccharide (LPS)-challenge model was approved by the Local Ethical Committee for Animal Experiments in Bydgoszcz, Poland under license numbers 27/2019 and 38/2021. The C57BL/6 male mice were housed in standard cages with an enriched environment under 12-hour day and night cycles. The air in the room was exchanged 15 times per hour, and temperature and humidity were maintained at 20-23°C and 50-60%, respectively. The animals had unrestricted access to food and water.

The TDAG8 (GPR65) double (−/−) knock-out (KO) C57BL/6 mouse strain was generated and purchased from Deltagen, Inc. (San Mateo, CA, USA). The KO and littermate control mouse breeding was approved by the Veterinary Authority of the canton of Zurich (in-house registration number 100635). All the animals were housed in a specific pathogen-free (SPF) facility. The animals were kept in type II long clear-transparent individually ventilated cages (IVCs, Allentown, New Jersey, USA). They were fed a pelleted and extruded mouse diet (R/M–H Extrudat, Ssniff Spezialdiäten, Soest, Germany) *ad libitum*. The light/dark cycle in the room was given through natural daylight. The temperature was set to 21 ± 1°C, with a relative humidity of 55 ± 5% and 75 complete changes of filtered air per hour.

### Mouse Brain Isolation and Immunohistochemistry

After decapitation whole brains from 20 weeks old male TDAG8 KO and wild-type (WT) mice were collected, carefully separated into hemispheres, and snap-frozen in liquid nitrogen. The hemispheres used for RT-qPCR were ground using mortar and pestle, homogenized in Fenozol reagent (A&A Biotechnology, 203-50), and stored at -20°C. For immunohistochemistry, the hemispheres were embedded in the O.C.T. matrix (VWR, 00411243). The tissue O.C.T. blocks were sectioned into 10 μm sagittal slices and mounted onto microscope slides. The slices were fixed in 4% ice-cold paraformaldehyde for 30 min at room temperature (RT) and washed with PBS. To avoid unspecific binding the slices were incubated in a blocking solution (10% BSA, 0.5% Triton X, 1% NGS in PBS) for 24 hours (h). Then the primary antibody solution (2% BSA, and 0,1% Triton-X in PBS) was applied and slices were incubated overnight in a humid box. All primary antibodies used in this study are shown in **Supplementary Table 1**. The slides were washed 3x for 30 min in PBS and incubated in a secondary antibody solution containing 1:500 dilution of anti-rabbit Cy3-conjugated (Jackson ImmunoResearch, 111-165-144) and anti-mouse Cy-5 conjugated (Jackson ImmunoResearch, 515-605-003) antibodies for 24 h. After mounting on glass cover slides the slices were imaged with Zeiss LSM880 (Zeiss, Oberkochen, Germany) confocal microscope and analysed using Zeiss Zen 3.5 software.

### LPS Challenge Model

Male C57BL/6 wild-type mice were used in the model. The mice received intraperitoneal (i.p.) injections of 0.9% NaCl (vehicle) or 2 mg/kg LPS for 24 h. The mice were anaesthetized with isoflurane and perfused transcardially with NaCl. Whole brains were removed and the right hemisphere was snap-frozen in liquid nitrogen. The hemispheres were homogenized with a pestle and a mortar in liquid nitrogen. The resulting powder was then suspended in Fenozol reagent (A&A biotechnology, 203-50). Total RNA was isolated from approximately 50 mg of homogenized tissue using the Total RNA Mini kit (A&A Biotechnology, 031-100).

### Cell Culture and Differentiation

MO3.13 human oligodendrocytic hybrid cell line (RRID: CVCL_D357) was purchased from Tebu-Bio (2018, CLU301, batch 131117 P25) and cultured in high glucose DMEM (Sigma Aldrich, D5796), 10% FBS and 1% pen/strep. Cells were plated on 6-well cell culture-treated plates and differentiated using 100 nM phorbol 12-myristate 13-acetate (PMA) (Sigma-Aldrich, P1585) for 72 h in serum-free RPMI 1640 medium (SFM, Sigma Aldrich, R8758) supplemented with 2 mM Glutamax and 20 mM HEPES with pH adjusted to pH=7.6 (high pH), pH=7.25 (normal pH) or pH=6.8 (low pH). The medium in each group was changed every 24 h. After treatments, MO3.13 cells were washed in PBS and scraped in 200 μl of Fenozol reagent (A&A biotechnology, 203-50). Total RNA was isolated using the Total RNA Mini kit (A&A Biotechnology, 031-100).

### pH Shift Experiments

The pH shift experiments and differentiation experiments were conducted in SFM supplemented with 2 mM Glutamax and 20 mM HEPES. The pH of the solutions was adjusted using calibrated pH meter (Mettler Toledo, Seven Compact S220) with 1M NaOH or 1M HCL. The initial pH of the media was adjusted to pH=8.1 (high pH) or pH=6.1 (low pH). The media were filtered and equilibrated in a 5% CO_2_ incubator for at least 72 h before experiments, and the pH was measured again. After equilibration, the final pHs were 7.6 (high pH) and 6.8 (low pH), measured at RT. The pH of the normal, unadjusted SFM was 7.2. Undifferentiated MO3.13 cells were plated on 6-well cell culture-treated plates and grown until reached 70% confluence. The cells were starved for 4 h in a serum-free high pH medium. The high pH medium was removed and the pH shift was performed by adding the high, normal, or low pH medium. The cells were cultured for up to 72 h depending on the experiment. After treatments, the cells were washed in PBS and scraped in 200 μl of Fenozol reagent (A&A biotechnology, 203-50). Total RNA was isolated using the Total RNA Mini kit (A&A Biotechnology, 031-100).

### pH Shift Experiments In Inflammatory Conditions

The MO3.13 oligodendrocytes were plated on 6-well cell culture-treated plates and cultured until approximately 70% confluency. The cells were starved for 4 h in serum-free high pH medium and treated with LPS (100 ng/ml) (Sigma Aldrich, L4391) in serum-free high, normal, or low pH medium for 2 h or 24 h. For the IL17/TNFα challenge experiments, the cells were starved in a serum-free high pH medium and then incubated overnight with a cocktail of 10 ng/ml TNFα (R&D Systems, 210-TA-100/CF) and 50 ng/ml IL-17A (R&D Systems, 317-ILB-050) recombinant cytokines. After treatments, MO3.13 cells were washed in PBS and scraped in 200 μl of Fenozol reagent (A&A biotechnology, 203-50). Total RNA was isolated using the Total RNA Mini kit (A&A Biotechnology, 031-100).

### Cell Migration Assay

Undifferentiated MO3.13 cells were starved for 2 h before the experiment in serum-free high pH medium. Migration assay was performed using 8.0 μm cell culture inserts (Corning, 353097) on 24-well plates. Each treatment condition was duplicated and each experiment was repeated at least three times. The cells were trypsinized, centrifuged, and resuspended in a high pH medium to obtain 800.000 cells/ml solution. Subsequently, 400 μl of high pH medium, low pH medium, or high pH medium supplemented with 0,01 μM 7-alpha,25-dihydroxycholesterol (7α,25OHC) (Sigma Aldrich, 64907-22-8) was added to the bottom of the chamber. Subsequently, 80 μl of the cell solution was added carefully into the insert top chamber. After 30 min at RT, the top chamber was topped up with 80 μl of high pH medium. The cells were left to migrate to the lower chamber for 24 h. Then the media was discarded from the chamber and the remaining cells were removed with a cotton swab. The transwells were immersed in 250 μL of crystal violet staining solution (Sigma Aldrich, CAS 548-62-9) for 10 min at RT. Then, they were carefully dipped in a beaker with distilled water and left to air dry. When dried, images were taken with a light microscope, and the transwells were incubated in 250 μL of extraction solution (Methanol, Cell Biolabs, CBA-100) for 20 min on a shaker. The solution was transferred to a 96-well plate to read absorbance at 590 nm with a VICTOR NivoTM plate reader (Perkin Elmer, USA).

### Real-Time Quantitative Polymerase Chain Reaction (RT-qPCR)

For reverse cDNA transcription, the Transcriba reverse transcription kit was used (A&A Biotechnology, 4000-100) following the manufacturer’s instructions. The RT-qPCR was performed using Sensitive RT HS-PCR Mix (A&A Biotechnology,2017-149 2000) on the Light Cycler 480 (Roche) under the following conditions: 120 s at 50°C, the 20s at 95°C then 40 cycles at 95°C for 3 s and 60°C for 30 s. Total RNA isolated from cerebellar slices, TDAG KO and control WT mice was reverse transcribed using the High-Capacity cDNA Reverse Transcription Kit (Applied Biosystems, 4368814). RT - qPCR was performed on the LightCycler 480 (Roche) using the TaqMan FAST Universal Mastermix (Applied Biosystems, 4304437). Cycling conditions were: 20 seconds (s) at 95°C, then 45 cycles at 95°C for 3 s, and 60°C for 30 s. The FAM dye-labelled TaqMan probes (Applied Biosystems) were used in all experiments and are listed in **Supplementary Table 2**. The relative mRNA expression was determined using the ΔΔCt method, calculated from absolute quantification after normalization to the endogenous housekeeping gene (β-actin).

### Cell Immunohistochemistry

The MO3.13 oligodendrocytes were cultured on 8-well imaging slides (Millipore, PEZGS0816) covered with Poly-D-Lysine in SFM low pH, normal pH, or high pH medium for 24 h. The cells were washed once with cold PBS, fixed for 10 min in ice-cold 4% PFA, and permeabilized for 60 s in cold 100% methanol. Cells were incubated in blocking solution (0.5% NGS, 1% BSA, 0.1% tween-20 in PBS) for 60 min and then overnight in primary antibodies diluted in PBS containing 0.5% BSA and 0.05% tween-20 at 4°C. The cells were washed 3x with PBS and incubated for 1 h at RT in a secondary antibody solution containing 1:500 dilution of anti-rabbit Cy3-conjugated (Jackson ImmunoResearch, 111-165-144) and anti-mouse Cy-5 conjugated (Jackson ImmunoResearch, 515-605-003) antibodies. In the final step, the nuclei of the cells were counterstained with Hoechst dye (ThermoFisher, H1399), washed, and imaged with Zeiss LSM880 (Zeiss, Germany) confocal microscope and analysed using Zeiss Zen 3.5 software.

### Statistical Analysis

All Statistical analyses were performed using GraphPad Prism statistical software (version 8.0 or higher, RRID: SCR_002798). Multiple comparisons between the experimental groups were made using one-way analysis of variance (ANOVA) followed by Sidak’s post-hoc tests for comparisons of differences between selected experimental groups, Dunnett’s post-hoc tests for comparisons of the control with other means and Tukey’s post-hoc test for comparisons of all means with every other mean. Multiple comparisons between the experimental groups and normalised control were made using one sample t-test. Comparisons between two groups were made using independent student *t*-tests. All data are presented as means ± standard error of the mean (SEM). Each experiment was independently repeated at least 3 times (N=3). Throughout this article significant differences are indicated by asterisks: \**p* ≤ 0.05, \*\**p* ≤ 0.01, \*\*\**p* ≤ 0.001, \*\*\*\**p* ≤ 0.0001.

## RESULTS

### TDAG8 is upregulated in MS plaques

Large-scale GWAS studies identified TDAG8 as a risk gene for MS [13,14]. We, therefore, examined TDAG8 expression in the WM in MS and control human brains. The mRNA transcripts of TDAG8 were somewhat elevated in the normal-appearing WM in the MS brains compared to the matching WM areas in the control brains (no significance was reached). However, TDAG8 was significantly upregulated in the WM lesions (MS plaques) compared to the WM in the control brains **(Fig 1A)** suggesting that TDAG8 may play some role in the pathophysiology of MS. This data prompted us to examine if TDAG8 signalling is required for normal myelination in the mouse brain. To answer this question, we investigated OPC and oligodendrocyte markers at gene and protein levels in adult WT and TDAG8 KO mice brains. The data showed no differences between the WT and TDAG8 KO brains in the mRNA levels of the OPC marker PDGFRα, the early myelination marker CNPase, nor in the mature myelin marker MBP **(Fig. 1B)**. The immunohistochemical staining of the brain sections from the WT and TDAG8 KO mice also did not reveal qualitative differences in the PDGFRα, CNPase and MBP proteins **(Fig. 1C)** indicating that TDAG8 signalling does not play a role in myelination under normal physiological conditions *in vivo*.

**Figure 1.**
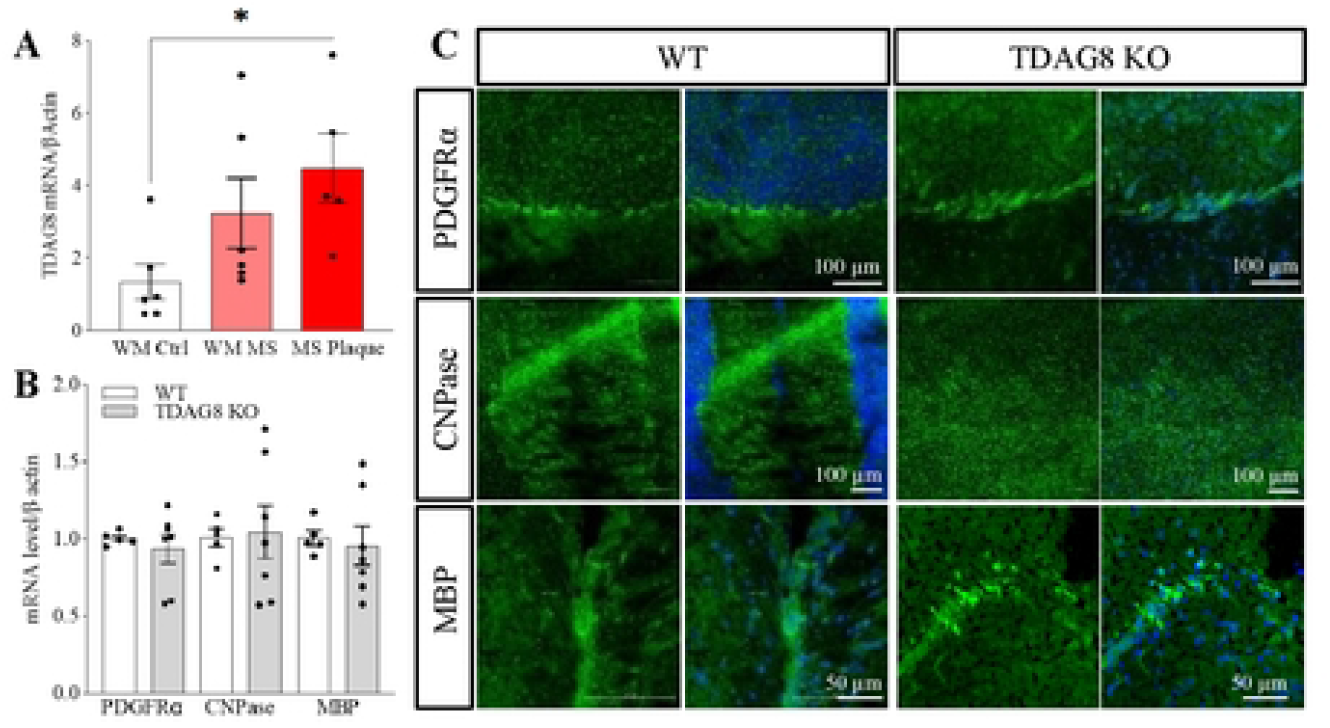
TDAG8 is upregulated in MS plaques. **A**. The mRNA expression of TDAG8 in normal-appearing WM in MS brains was slightly upregulated (241% +/− 72% vs. WM Ctrl. P>0.05> and significantly upregulated in demyelinating plaques (335% +/-71% vs. WM Ctrl). One-way ANOVA and Dunnett’s post-hoc tests. *p≤0.05. N=5-6 human brains. **B.** The analysis of oligodendrocyte and myelin markers in whole brain homogenatcs of TDAG8 KO mice did not reveal differences between the KO and WT mice. Student t-tests. p>0.05. N=5 WT and N=7 TDAG8 KO mice. **C.** Representative images of inununohistocheinically stained WT and TDAG8 KO mice brains show no differences in myelination between the two genotypes.

### TDAG8 is upregulated in oligodendrocytes during maturation and under acidic conditions

Neuroinflammation, neurodegeneration or ischemic stroke are associated with local tissue acidification [10,25–28]. Whether the pH-sensing receptor TDAG8 is involved in sensing acidic pH by oligodendrocytes has not been investigated. Here, we first examined TDAG8 expression in undifferentiated MO3.13 oligodendrocytes and during maturation. The data showed a steady increase in TDAG8 expression during MO3.13 maturation with significant upregulation of TDAG8 after 72 hours of stimulation with the maturation-inducing agent, PMA **(Fig. 2A)**. To examine if the expression of TDAG8 is pH-dependent we then performed pH shift experiments. A shift to low pH, when TDAG8 is active, upregulated its expression, but only temporarily (∼2 hours) and returned to baseline thereafter (∼12 hours) **(Fig 2B)**. While a shift to physiologically neutral pH significantly downregulated TDAG8 expression for at least 24 hours. Subsequently, we performed the pH shift experiments during oligodendrocyte maturation (PMA treatment) to observe the effects of pH on oligodendrocyte differentiation. The data showed no additional pH-mediated effect on PMA-induced TDAG8 expression at 72 hours (corresponding PMA-only treatment in Fig 1A, the 72-hour timepoint) **(Fig. 2C)**. However, analysis of the oligodendrocyte differentiation markers revealed that the low pH maintained the expression of the OPC marker PDGFRα, which is normally downregulated as MO3.13 oligodendrocytes mature [29] **(Fig. 2D)**. The low pH also potentiated the expression of the mature oligodendrocyte marker, myelin basic protein (MBP), which increases as MO3.13 oligodendrocytes mature [29]. We then examined the morphology of oligodendrocytes after shifting to normal (pH ∼7.3), high (pH ∼7.6) or low pH (∼6.5). The data showed no changes in oligodendrocyte morphology when shifting to normal or high pH (silencing the receptor) **(Fig. 2E and F)**. However, a shift to low pH, induced oligodendrocyte branching. No qualitative differences in the level of PDFRGα, CNPase or A2B5 in oligodendrocytes were observed after 24 h culture in either normal, high or low pH.

**Figure 2.**
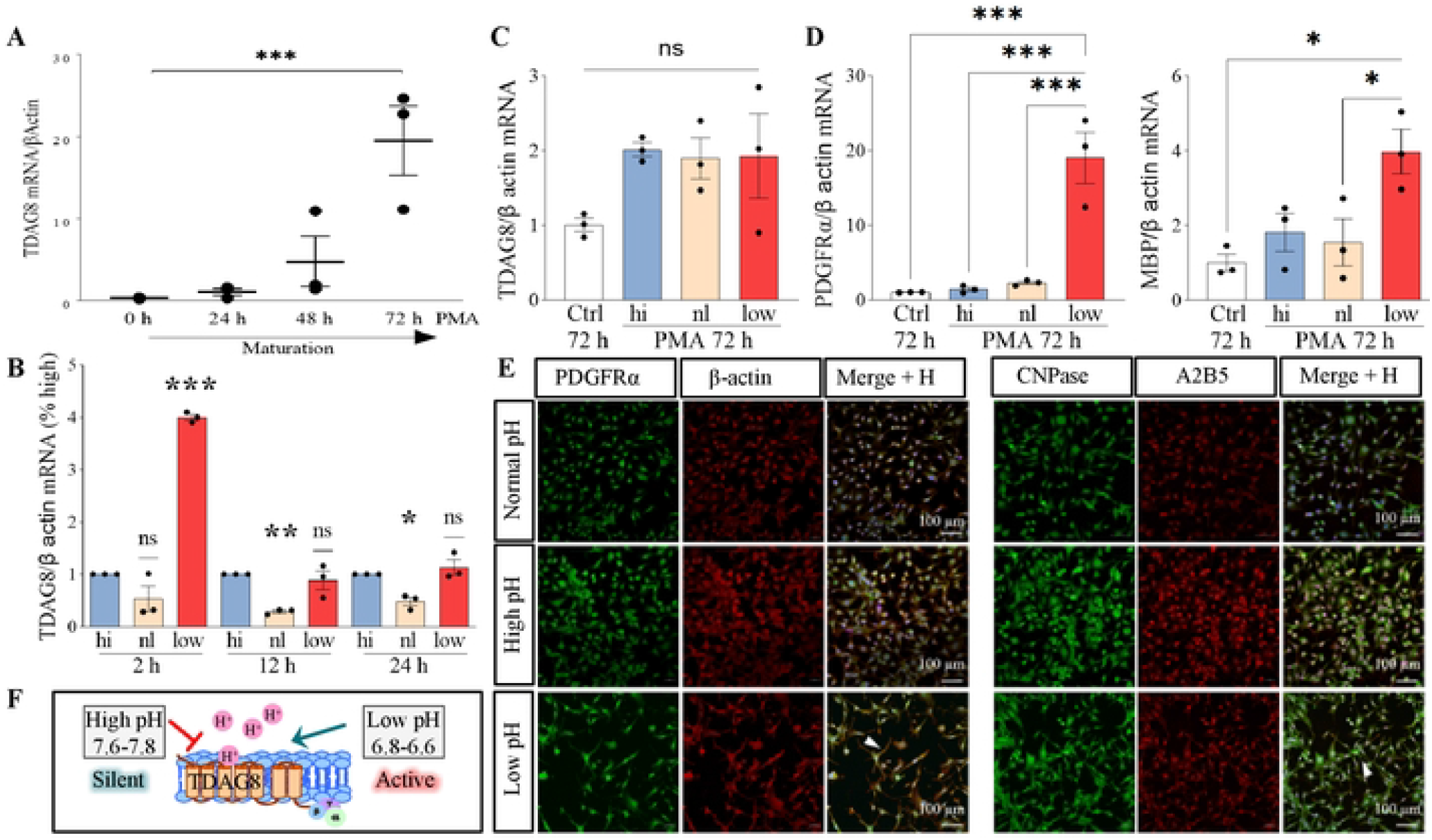
TDAG8 is upregulated in oligodendrocytes during maturation and under acidic conditions. **A**. The mRNA expression of TDAG8 steadily increases in PMA-stimulalcd MOT 13 oligodendrocytes (60.38%+/− 13.11%0 h vs. 72 h). One-way ANOVA and Dunnett’s post-hoc tests. ***p≤0.00]. N=3 independent experiments. **B**. Low pH (6.6-6.8) strongly induces TDAG8 expression after 2 h (400% +/-6% vs. Hi) and dow nrcgulatcs it in normal pH after 12 h (29%+/−3% vs. Hi), One sample t-test. *p≤;0.05, **p≤0.0l. ***p≤0.001. N=3 independent experiments. **C**. A pH shift to cither low or high pH during PMA -induced maturation of MO3.13 oligodendrocytes docs not aflcct TDAG8 expression. One-way ANOVA and Tukcy’s post-hoc tests. p>0.O5. N=3 independent experiments. **D**. The PDGFRα mRNA transcripts arc maintained in low pH after long-term (72 h) treatment (190% +/-34,4% vs. Ctrl) and MBP transcripts arc uprcgulatcd (39.7% +/-9.7% vs. Ctrl). One-way ANOVA and Tukcy’s post-hoc tests. *p≤0.05 ***p≤0.001. N=3 independent experiments. **E**. Representative images showing MO3.13 oligodendrocy tes stained with anti-PDGFRα (green). anti-CNPasc (green) and anti-A2B5 (red) antibodies after 24-It our culture in cither normal, high or low pH. The low pH-induccd elongation of cell processes (arrows). Normal pH=7.2; high pH=7.6. low pH=6.8. **F**. Schematic representation of TDAG8 signalling in different pH.

### Acidic pH inhibits oligodendrocyte chemotaxis

OPCs are highly motile cells. An ability crucial to their function especially in the context of remyelination when new myelin is formed from surviving adult oligodendrocytes or by recruited OPCs attracted to the site of injury by various chemoattractants and acidic pH [24]. Because in our experiments acidic pH induced the expression of TDAG8 and maintained the expression of OPC marker PDGFRα, we investigated the effects of TDAG8 activation (a shift to acidic pH) on the migration of undifferentiated MO3.13 oligodendrocytes. The data showed that a pH shift from high (silent receptor) to low (receptor maximally active) had a minor chemotaxis-inducing effect on oligodendrocytes compared to no pH shift (maintenance in high pH) **(Fig. 3)**. Subsequently, to compare the level of migration induced by low pH to a known chemotaxis-inducing agent in MO3.13 oligodendrocytes and other glial cells [29,30], we stimulated the cells with EBI2 receptor agonist, oxysterol 7α,25OHC, in either low or high pH. The MO3.13 oligodendrocytes extensively migrated towards 7α,25OHC in high pH. Strikingly, the 7α,25OHC-induced chemotaxis was significantly reduced in low pH indicating that an acidic environment inhibits oligodendrocyte migration. The data thus far presented on the role of TDAG8 in oligodendrocytes indicates that TDAG8 may be involved, to some extent, in oligodendrocyte differentiation under acidic conditions and that low pH inhibits oligodendrocyte migration *in vitro*.

**Figure 3.**
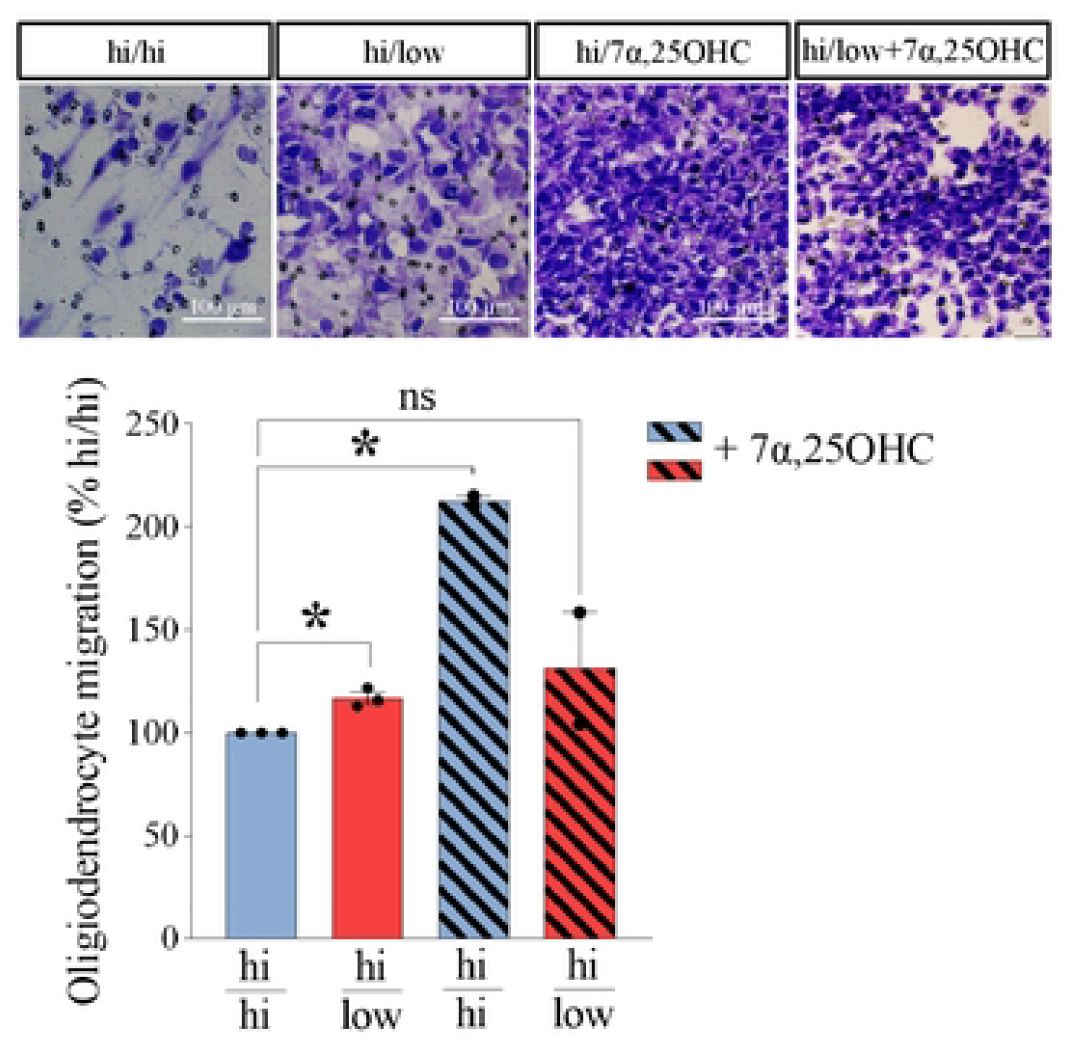
Acidic pH inhibits oligodendrocytes chemotaxis. A pH shift from high to low pH stimulates the MO3.13 oligodendrocytes chemotaxis (117% +7-2,6% vs. hi/hi). The EBI2 receptor agonist 7α25OHC in hi pH medium induces an extensive cell migration (212% +/ - 2.5% vs. hi/hi). However. 7α.25OHC-induccd chemotaxis is inhibited in low pH (131.5% +/-27.2% vs. hi/hi) One sample t-test, *p≤0.05. N=3 independent experiments.

### TDAG8 is upregulated in the brain during systemic inflammation

I.p. injections of LPS were shown to induce neuroinflammation and significant pH reduction in the mouse brain [26,31]. Moreover, the pH-sensing receptor, TDAG8, is mainly expressed in the immune cells and has anti-inflammatory properties [32–35]. Because our investigations into the role of TDAG8 in oligodendrocytes thus far indicated that TDAG8 is upregulated in oligodendrocytes in an acidic environment (pH 6.8), which is when TDAG8 is maximally activated, we asked the question of whether TDAG8 is involved in inflammatory signalling in the brain. We first induced inflammation in cultured MO3.13 oligodendrocytes during a pH shift with a cocktail of TNFα/IL17 cytokines, which were shown before to induce pro-inflammatory signalling and cytokine release in glial cells [31,36]. Here, in the human MO3.13 oligodendrocytes, the overall TDAG8 expression was downregulated after 18 hours of TNFα/IL17-treatment in high, normal and low pH with significance reached for cells grown in high and low pH **(Fig. 4A)**. We then tested the effects of bacterial LPS on TDAG8 expression in these cells *in vitro*. The data showed that stimulation of oligodendrocytes with LPS significantly induces mRNA expression of TDAG8 short-(2 h) and long-term (24 h) **(Fig. 4B)**. Subsequently, we injected mice with either LPS or vehicle and examined TDAG8 expression in whole brain homogenates. Similarly to the *in vitro* experiments, i.p. injection of LPS strongly induced TDAG8 expression in the mouse brain after 12 and 24 hours **(Fig. 4C)** implicating that TDAG8 may be involved in the regulation of pro-inflammatory signalling in the CNS and specifically in oligodendrocytes.

**Figure 4.**
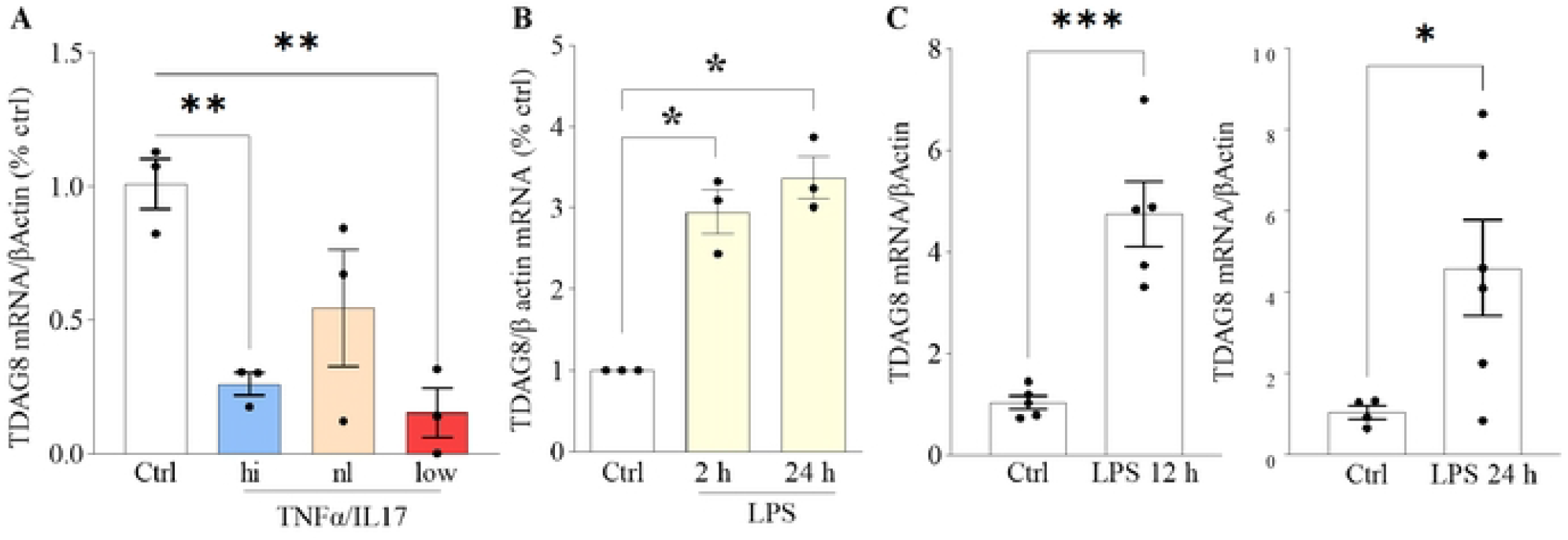
TDAG8 is upregulated in the mouse brain during systemic inflammation. **A**. A combined treatment of MO3.13 oligodendrocytes with a cocktail of TNFα/IL17 pro-inflammatory cytokines downrcgulalcd TDAG8 expression irrespective of the pH (15% +/ - 9% hi vs. Ctrl and 26% +/ - 4% low vs. Ctrl). One-way ANOVA and Dunnett’s post-hoc tests **p≤0.01. N=3 independent experiments. **B**. Conversely, treatment with bacterial LPS (100 ng/ml) for 2 h (295% +/- 27% vs. Ctrl) or 24 h (337%+/- 26% vs. Ctrl) induced TDAG8 expression. One sample t-test. *p≤0.05. N=3 independent experiments. **C**. I.p. LPS injections strongly upregulalc TDAG8 expression in the mouse brain after 12 h (460% +/ - 62% vs. vehicle) and 24 h (441% +/-114% vs. vehicle). Student t-tests. *p≤0.05 * * *p≤0.001. N=4-6 mice per condition.

## DISCUSSION

Here we explored the expression and function of the pH-sensing receptor TDAG8 in human oligodendrocytes and inflamed CNS. We found that TDAG8 is steadily upregulated during oligodendrocyte maturation and, temporarily, in low pH. Interestingly, the data revealed morphological changes (elongated processes) in oligodendrocytes, also only at low pH. However, there were no differences in normal myelination in the TDAG8 KO mice brains, nor did the low pH induce oligodendrocyte chemotaxis *in vitro*. Conversely, bacterial LPS strongly upregulated TDAG8 in oligodendrocytes *in vitro* and mouse brains *in vivo*. Our explorations culminated in experiments using human material from MS patients where we found that TDAG8 is upregulated in MS plaques.

Differentiation of the MO3.13 oligodendrocytes *in vitro* can be achieved by stimulation with either PMA or by culturing in SFM [37,38]. The cell maturation is indicated by reduced expression of the OPC or pre-oligodendrocyte markers and an increase in mature and myelinating oligodendrocyte markers [29,39]. Here, we used both, the PMA and SFM (ctrl) to induce the differentiation of MO3.13 oligodendrocytes. Our data demonstrated that TDAG8 transcripts steadily increase during PMA-induced maturation and only temporarily in low pH SFM. Interestingly, only when combined, the low pH and PMA sustained the expression of the OPC marker PDGFRα and induced the expression of mature oligodendrocyte marker MBP suggesting that TDAG8 may be involved in OPC/pre-oligodendrocyte function related to myelination under pathophysiological conditions (low pH). Indeed, examination of the cell morphology revealed elongation of the cell processes only in the acidic pH. A previous study using primary rat oligodendrocytes also found that acidic pH induces elongation of cell processes but inhibits cell differentiation, as indicated by low MBP levels [24]. These contrasting observations may result from different experimental conditions of PMA/low pH for 3 days in our study versus low pH for 5 days in Jagielska et al. (2013). Bernard and colleagues (2006), on the other hand, observed the highest differentiation of primary rat cerebellar oligodendrocytes at pH 7.15 and a declining rate of differentiation (GalC levels) at either lower or higher pH ranges. None of the previous studies examined the OPC marker PDGFRα, which was strongly upregulated (or sustained) in our study in the low pH.

Fetal human forebrain-derived OPCs enriched by PDGFRα (CD140α) are most effective in both, myelinating and migratory capacities, compared with those selected by A2B5, when transplanted to hypomyelinated shiverer mouse brain, suggesting that myelin disorders may be promising targets for cell-based therapy [41,42]. Whether acidic pH indeed inhibits oligodendrocyte maturation and/or maintains OPCs in an undifferentiated state (PDGFRα expression), and whether TDAG8 is involved in these processes needs to be investigated further. Low pH could act on other receptors that have been shown to regulate OPC differentiation and migration, such as EBI2 or the mechanosensing ion channel Piezo1, which is inhibited in pH under 6.9 [29,38,43,44]. Nevertheless, looking at myelination *in vivo* in untreated TDAG8 KO mice, our explorations demonstrate that TDAG8-mediated signalling is not required for normal myelination in the mouse CNS.

Local changes in pH, which accompany inflammation and tissue injury, should attract local and peripheral immune cells, as well as OPCs, to the affected area to effectively prepare the milieu for regeneration and remyelination. Our investigations indicated that a shift from high to low pH moderately induced oligodendrocyte chemotaxis and inhibited 7α,25OHC-induced migration indicating that low pH has an overall inhibitory effect on oligodendrocyte chemotaxis. Similar observations were made by Jagielska and colleagues (2013) who found that at a fixed low pH of 6.0, the migration velocity and radius of primary rat oligodendrocytes are lower compared to the pH ranges of 6.5 and higher (pH 6.8 in our experiments). However, when a pH gradient was formed in Zigmond chambers the oligodendrocytes migrated towards the acidic pH (pH 6 versus pH 7) indicating that OPCs may indeed be attracted to the lesion or inflammatory site where a pH gradient forms, with more acidic pH closer to the wound. Neither of the previous research discussed here, including ours, used TDAG8 ligands rendering it impossible to conclude whether TDAG8 is involved in oligodendrocyte migration in acidic pH. In light of the lack of well-characterised, potent and selective TDAG8 ligands (either inhibitors or agonists), future investigations should use TDAG8-deficient oligodendrocytes to decipher whether TDAG8 modulates oligodendrocyte migration.

The pH-sensing receptor, TDAG8, is mainly expressed in the immune cells and modulates inflammatory signalling in various diseases (reviewed in Sisignano et al., 2021). We, therefore, investigated whether an inflammatory challenge of either cells *in vitro* or mice *in vivo* with different pro-inflammatory agents affects TDAG8 expression in the brain. Previous studies showed that a combined treatment of cells *in vitro* with TNFα and IL17 induces an extensive release of pro-inflammatory cytokines including IL6 involving the NFκB signalling pathway [36,46]. Importantly, the release of the pro-inflammatory mediators is greater when IL17 and TNFα are combined than when either is given alone. Specifically in oligodendrocytes, treatment with IL17 was shown to synergistically augment the TNFα-induced oligodendrocyte apoptosis, mitochondrial dysfunction, release of reactive oxygen species and induce an arrest of OPC maturation, thus promoting myelin damage [46]. In our experiments, the co-treatment of MO3.13 oligodendrocytes induced an overall downregulation of TDAG8 transcripts irrespective of the extracellular pH but most pronounced in low and high pH respectively. Therefore, our results demonstrate that the anti-inflammatory signalling through TDAG8 receptor is directly affected by the cytokines implicated in MS pathogenesis [47–51]. However, the non-pH-specific reaction to TNFα/IL17 renders it difficult to draw any conclusions regarding the possible role of TDAG8 in the TNFα/IL17-mediated oligodendrocyte dysfunction and should be dissected in future studies using receptor ligands or TDAG8 KO animals. A comprehensive investigation could deliver a novel mechanism for limiting oligodendrocyte loss in demyelinating diseases such as MS.

The increased TDAG8 expression reported here after LPS-challenge in cultured oligodendrocytes and the brain is in line with the previous reports of the anti-inflammatory role of TDAG8 in LPS-stimulates peritoneal macrophages and T cells [52,53]. In these studies activation of TDAG8 in acidic pH reduced the levels of pro-inflammatory cytokines (TNFα, IL6) and increased the synthesis of the anti-inflammatory IL10 cytokine after LPS challenge. In the LPS-induced acute lung injury model, the expression of TDAG8 was also induced in the lungs and local macrophages, and the lung damage was increased in TDAG8-deficient mice, again indicating the anti-inflammatory role of TDAG8 [54]. Our investigations suggest that TDAG8 is involved in inflammatory signalling in the brain upon peripheral bacterial infection and should be further elucidated to answer such questions as to whether modulation of its signalling may be used to attenuate neuroinflammation and associated cell injury and demyelination.

As for the increased TDAG8 transcripts in the WM plaques, we believe that these are most likely due to increased infiltration by reactive lymphocytes, which highly express TDAG8, and are enriched in MS plaques [55]. An RNA sequencing study found a decreased expression of TDAG8 in the corpus callosum and the optic chiasm of MS patients [56]. These results indirectly support our conclusions that the source of the increased TDAG8 expression in the plaque are the invading lymphocytes. Whether TDAG8 is upregulated also in the CNS and immune competent resident cells such as microglia or astrocytes in the MS brain should, however, be investigated to possibly open new opportunities to modulate glia reactivity and inflammatory signalling in MS.

Our explorations of the expression and function of TDAG8 in the CNS and oligodendrocytes here reported provided several interesting routes for future investigations. Specifically, the neuroinflammatory line of inquiry including the potential role of TDAG8 in the LPS and TNFα/IL17-mediated oligodendrocyte function and the role of TDAG8 in the resident glial cells, in particular astrocytes and microglia, in the pathophysiology of MS. Closely related to MS is also the question whether TDAG8 modulates OPC differentiation and interaction between OPCs/oligodendrocytes and immune cells under pathophysiological conditions, that is tissue acidosis, that accompanies neuroinflammation, demyelination and neurodegeneration.

## STATEMENTS

### Author Contribution

C.V., K.S., G.R., B.K., A.R., conceived the study or provided merit-based support; M.O., F.C., M.V.E., C.V., A.R. established the methodology; M.O., F.C., M.H., M.V.E., A.R. performed the experiments; M.O., F.C., A.R. analysed the data; A.R. visualised the data; M.O., A.R. wrote the original manuscript; A.R. supervised the project; G.R., A.R., acquired funding; B.K., A.R. provided project administration and coordination; All authors read, reviewed and approved the manuscript.

### Data availability

The data supporting the conclusions of this article will be made available by the authors, without undue reservation, to any qualified researcher.

### Funding

This project received funding from the National Science Centre, Poland, grant registration number: 2019/33/B/NZ4/03000 (AR) and the Swiss National Science Foundation, grant numbers: 310030_172870, 314730_153380 (GR).

### Conflict of Interest Disclosure

The authors declare no conflicts of interest

### Ethics Approval And Patient Consent Statement

Written informed consent was obtained from all human participants in accordance with the Declaration of Helsinki, and the study was approved by the Independent Bioethics Committee For Scientific Research at Medical University of Gdańsk, Poland under the licence number: NKBBN/457/2019.

Animal experiments were approved by the Local Ethical Committee for Animal Experiments in Bydgoszcz, Poland under licence numbers 27/2019 and 38/2021 and the Veterinary Authority of the canton of Zurich, Switzerland (in-house registration number 100635).

## ACKNOWLEDGMENTS

Human brain specimens from MS patients and controls were obtained from the Rocky Mountain MS Center Tissue Bank (Englewood, United States). We acknowledge the support of the National MS Society to the Rocky Mountain MS Center Tissue Bank for providing the human specimens.

We would like to thank Dr. Ilona Klejbor and Ms. Beata Muszyńska for their support with the LPS mouse model.

## Notes

### Competing Interest Statement

The authors have declared no competing interest.

